# Widespread regulatory specificities between transcriptional corepressors and enhancers in *Drosophila*

**DOI:** 10.1101/2022.11.07.515017

**Authors:** Jelle Jacobs, Michaela Pagani, Christoph Wenzl, Alexander Stark

## Abstract

Animal development and homeostasis critically depend on the accurate regulation of gene transcription, which includes the silencing of genes that should not be expressed. Repression is mediated by a specific class of transcription factors (TFs) termed repressors that, via the recruitment of co-repressors (CoRs), can dominantly prevent transcription, even in the presence of activating cues. However, the relationship between specific CoRs and enhancers has remained unclear. Here, we used functional genomics to uncover regulatory specificities between CoRs and enhancers. We show that enhancers can typically be repressed by only a subset of CoRs. Enhancers classified by CoR sensitivity also show distinct biological functions and endogenous chromatin features. Moreover, enhancers that are sensitive or resistant to silencing by specific CoRs differ in TF motif content, and their sensitivity to CoRs can be predicted based on TF motif content. Finally, we identified and validated specific TF motifs that have a direct impact on enhancers sensitivity or resistance towards specific CoRs, using large scale motif mutagenesis and addition experiments.

This study reveals the existence of TF motif-based regulatory rules that coordinate CoRs-enhancer compatibilities. These specificities between repressors and activators not only suggest that repression occurs via distinct mechanisms, but also provide an additional layer in transcriptional regulation that allows for differential repression at close genomic distances and offers multiple ways for de-repression.

## Introduction

Animal development and homeostasis critically depend on differential gene expression(*1, 2*), enabled by the precise regulation of transcriptional activation and repression(*3*–*8*). Although repression is often associated with heterochromatin(*9, 10*), genes can also be silenced in transcriptionally permissive euchromatin(*11*–*14*) by repressive transcription factors (TFs), also termed repressors, that bind to DNA and recruit corepressors (CoRs). As CoRs can suppress transcription even in the presence of activators(*15, 16*), this mode of gene silencing is termed *active transcriptional repression*(*17, 18*). Active repression is critical and its failure can cause developmental defects(*19*–*21*) and diseases like cancer(*12, 22*); and it is conceptually intriguing as it requires the fast and efficient overriding of activating cues. However, the modes and mechanisms of this process are unclear, and whether a regulatory code coordinates repression and activation, is unknown.

Given that transcriptional activation can occur via distinct and mutually incompatible modes(*23*– *25*), it is intriguing to speculate whether distinct modes of active transcriptional repression exist. Examples of specificities between repressors and activators have indeed been observed. The CoR Retinoblastoma protein (Rb) as part of the DREAM complex for example can repress the TFs E2F, Mip120 and PU.1 but not others like SP-1(*26*–*28*), and repressors can have different activities in distinct transcriptional contexts(*29*). Comprehensive studies allowing to define different modes of active repression and uncover their regulatory rules are however lacking.

Here, we determined the mutual compatibilities between five known CoRs and a genome-wide library of active enhancers by measuring enhancer-activity changes upon CoR tethering in otherwise unperturbed cells, similar to activator-bypass experiments(*23, 30*). We reasoned that testing each enhancer with each CoR in all possible combinations should reveal CoR-enhancer combinations that lead to decreased enhancer activity (enhancers are *sensitive*) and those that do not (*resistant*), indicative of compatible and incompatible pairings.

## Results

### UAS-STARR-seq for genome-wide screening of enhancer-corepressor specificity

We first wanted to test whether distinct specificities between CoRs and enhancers exist, whereby certain enhancers are sensitive to repression by a given CoR, while other enhancers are resistant. For this, we need to systematically measure the effects that selected CoRs have on the activity of a large number of enhancers.

The comprehensive mapping of CoR-enhancer compatibilities, by examining all combinations of CoRs and enhancers, requires highly controllable quantitative high-throughput assays. We therefore modified the massively parallel enhancer-activity assay STARR-seq(*31*) to enable the function-based testing of genome-wide enhancer candidate libraries with different CoRs. Briefly, we introduced four upstream-activating-sequence (UAS) motifs immediately downstream of the enhancer library, which leave the enhancer sequence intact yet allows for the direct tethering of selected CoRs via the Gal4 DNA-binding domain (Fig 1A, see Methods). The tethering of CoRs next to active enhancers directly assesses whether CoRs can override existing activating cues, a process akin to active repression.

**Fig. 1.**
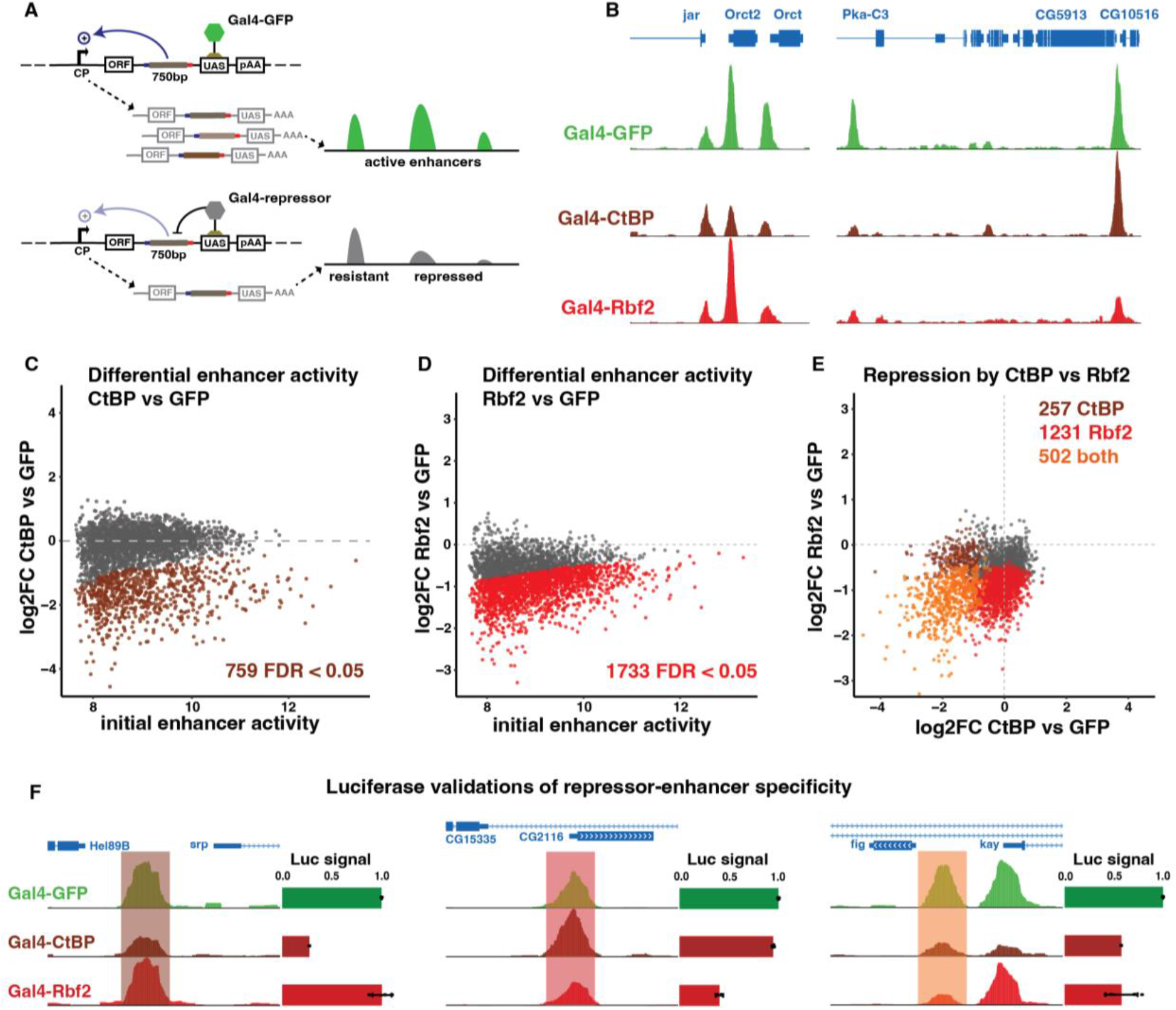
UAS-STARR-seq maps specificities between CoRs and active enhancers. (**A**) Schematic of the UAS-STARR-seq high-throughput enhancer-CoR assay. Gal4-GFP or a Gal4-CoR is recruited to the UAS sites near the genome-wide library of candidate enhancers. (**B**) UCSC screenshot visualizing the activity of nine enhancers (peaks) when GFP (green), the CoR CtBP (brown) and the CoR Rbf2 (red) are recruited. (**C-D**) MA-plots of initial enhancer activity (log2 normalized reads Gal4-GFP) on x-axis and repression by CtBP or Rbf2 respectively on y-axis, log2FC(enhancer activity Gal4-CoR vs Gal4-GFP), n=3094. (**E**) Scatterplot contrasting the sensitivity to repression (log2FC) of each enhancer (n=3094) towards CtBP and Rbf2. Orange for enhancers sensitive to both CoRs (FDR < 0.05 and log2FC < 0), brown for enhancer sensitive to CtBP and red for enhancers sensitive to Rbf2. (**F**) Luciferase validations, each time the tested enhancer is marked with a transparent column and the normalized luciferase activity is displayed as horizontal bar plots, n > =2.

We chose *Drosophila* S2 cells as a model system and a panel of five CoRs; CoRest, CtBP, Rbf, Rbf2 and Sin3A. These CoRs represent different protein complexes, repressive pathways, enzymatic functions, and distinct groups with context-specific functions(*16, 32*–*34*). Testing diverse CoRs casts a wide net and should increase our ability to detect compatible and incompatible CoR-enhancer pairs. For each CoR, we performed two independent UAS-STARR-seq screens where we co-transfect the UAS-STARR-seq library with a vector that expresses the Gal4-CoR (or Gal4-GFP as neutral control; Fig 1A) and spike-in-controls for normalization (see Methods). As spike-in controls we used a distinct STARR-seq library containing 18 *Drosophila pseudoobscura* enhancers, cloned without the UAS motifs, and hence not targeted by the Gal4-CoRs. In all cases, the two independent replicates correlated well (PCC > 0.84, fig. S1 A-F). In order to reliably assess repression (which requires a high baseline activity), we decided to evaluate enhancer-activity changes for 3094 enhancers that were highly active in Gal4-GFP controls.

### The CoRs CtBP and Rbf2 silence different subsets of enhancers

First, we determined which of the 3094 enhancers could be silenced by the highly conserved CoR CtBP(*35, 36*), by assessing the enhancer activity changes when tethering Gal4-CtBP versus Gal4-GFP. This revealed that some enhancers, like the enhancers near *Orct, Orct2* and *Pka-C3*, were reproducibly repressed by Gal4-CtBP, whereas others, like the enhancer near *CG10516*, were unaffected (Fig 1B). A differential analysis based on the two replicates per condition using edgeR (see methods) showed that CtBP significantly (FDR < 0.05, log2FC < 0) repressed 759 enhancers but not the remaining 2335 (Fig 1C). Thus, CtBP could only repress ∼25% of the enhancers, suggesting that CtBP displays preferences or specificities towards some enhancers but not others.

To test whether these specificities vary and depend on the recruited CoR, we next determined the enhancer-activity changes upon recruiting Rbf2, a CoR from the retinoblastoma protein family(*37*). Interestingly, Rbf2 was also able to repress only a subset of the enhancers (1733, 56%, Fig 1D), including the enhancers near *Orct* and *Pka-C3*, which were also repressed by CtBP, and *CG10516*, which was not repressed by CtBP. In contrast, it was unable to repress the aforementioned *Orct2* enhancer and others, indicating that the specificities of Rbf2 differ from those of CtBP (Fig 1B). Indeed, while 502 enhancers were repressed by CtBP and Rbf2, 1231 enhancers were repressed by Rbf2 but not by CtBP and, vice versa, 257 enhancers were repressed by CtBP but not by Rbf2 (Fig. 1E).

These enhancer-CoR specificities validated in luciferase reporter assays, in which the CoRs were recruited upstream of the enhancer and promoter (Fig 1F, see methods). This assay confirmed that an enhancer near *serpent* was specifically repressed by Gal4-CtBP but not Gal4-Rbf2, an enhancer near CG2116 was specifically repressed by Gal4-Rbf2 but not Gal4-CtBP, and an intronic enhancer of *kay* was repressed by both CoRs, each in agreement with the STARR-seq results (Fig. 1F). Taken together, the CoRs CtBP and Rbf2 are each able to repress a specific subset of enhancers but not others (Fig. 1B-F), indicating the existence of distinct CoR-enhancer specificities.

### Differential sensitivity to five CoRs define distinct types of enhancers

Screening three additional prominent CoRs; CoRest, Rbf and Sin3A revealed that each of them was able to repress a specific subset of enhancers (Fig. 2A, fig. S1 G-I). CoRest was the strongest repressor, repressing 1452 enhancers and often reducing their activity to background levels (fig. S1G). The overall repression profiles of CoRest and Sin3A were similar to that of CtBP (PCCs of 0.83 and 0.7 respectively) and clearly different from Rbf2 (PCC 0.28 and 0.24 respectively, fig. S1K). Rbf’s repression profile was not similar to any of the other tested CoRs (PCC < 0.05). In general, the five tested CoRs were each able to repress a subset of enhancers and displayed distinct specificities.

**Fig. 2.**
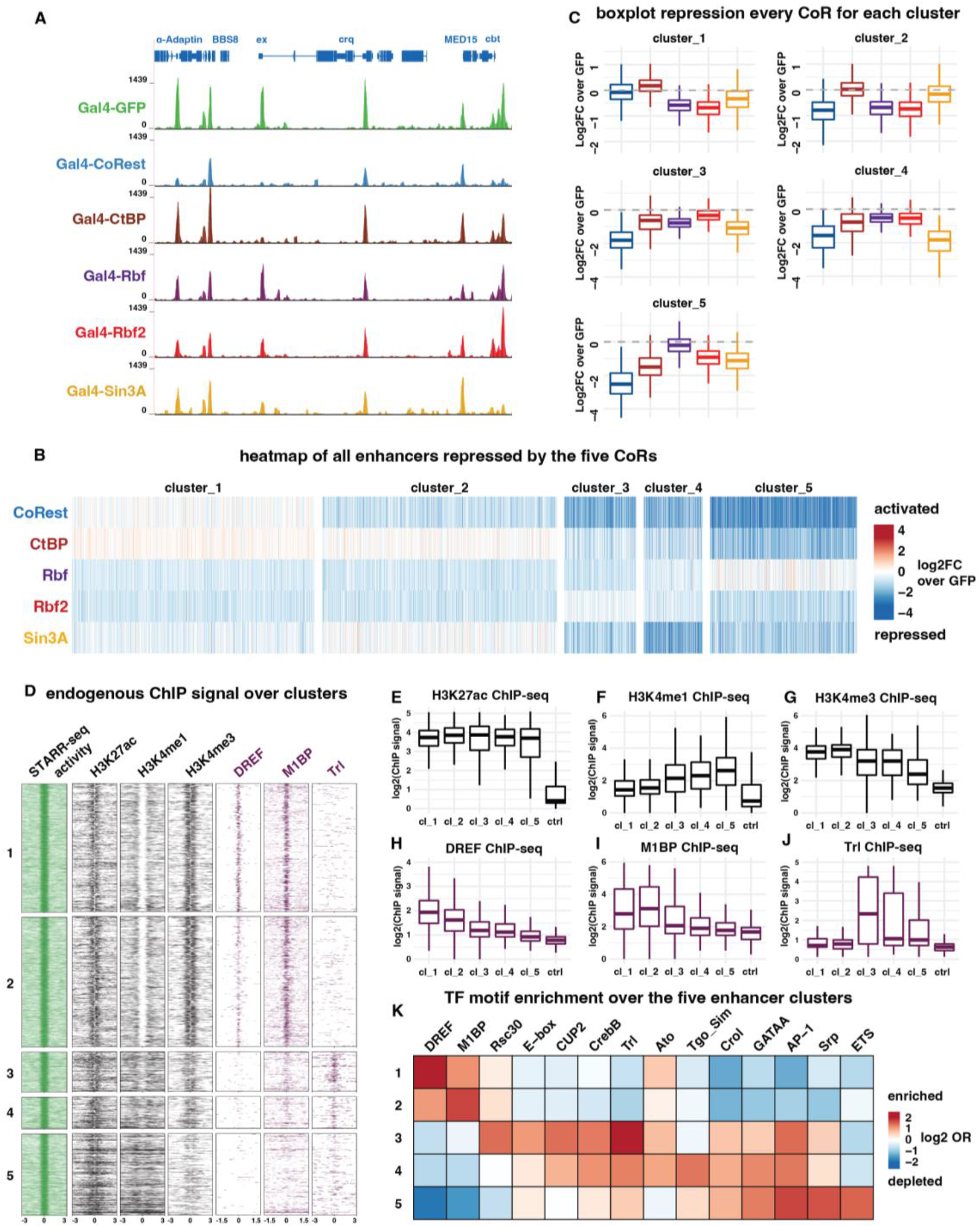
Enhancer clusters differ in chromatin marks, TF binding and sequence motifs. (**A**) UCSC screenshot of spike-in normalized UAS-STARR-seq tracks displaying the effect of each CoR on enhancer activity. (**B**) Heatmap of all enhancers from the genome-wide screen, grouped into 5 clusters based on their similarity in repression profile. The log2FC in enhancer activity is plotted between the gal4-CoR and gal4-GFP, repressed enhancers are marked in blue, unaffected in white and activated in red. (**C**) Boxplots summarizing the effect that each CoR has on the activity of enhancers from the 5 clusters. (**D**) Heatmaps visualizing endogenous ChIP-seq coverage over the enhancers in the different clusters. From left to right; STARR-seq activity under Gal4-GFP condition, chromatin marks H3K27ac, H3K4me1 and H3K4me3 and the TFs DREF, M1BP and Trl. (**E**-**J**) Boxplots summarizing the ChIP-seq signals over the different enhancer clusters, including negative control regions (1000 regions in the genome with no enhancer activity). (**K**) Heatmap of significantly enriched/depleted TF motifs log2 (odds ratio motifs in selected cluster vs all other clusters).

Given that enhancers were differentially sensitive to some of the tested CoRs, and resistant to others, we next explored whether these differential sensitivities were related to other enhancer properties. For this, we first clustered the enhancers into groups of similar repression patterns (Fig. 2B). We used a self-organizing tree algorithm (SOTA) with the PCC as distance metric to cluster the enhancers into five groups based on their sensitivity to the CoRs (Fig. 2B&C, log2-fold-change against Gal4-GFP, see methods). Cluster 1 contains CoRest and CtBP resistant enhancers while enhancers in cluster 2 are resistant to CtBP and Sin3A. Cluster 3 contains enhancers that are resistant to Rbf2, enhancers in cluster 4 are sensitive to all CoRs while enhancers in cluster 5 are overall very sensitive to repression but resistant to Rbf (Fig. 2C). Hence, the enhancers of S2 cells could be divided into five groups, defined by their differential response to the tested CoRs.

### Enhancers clustered by corepressor sensitivity differ in chromatin marks, transcription factor motif content and binding

Next, we investigated whether these enhancer clusters differ in additional properties that correlate with their behaviour towards the CoRs. Initial enhancer activity, as measured by UAS-STARR-seq with Gal4-GFP, was similar for all five clusters (Fig. 2D, fig. S3A). Also H3K27ac, a histone modification that marks active enhancers(*38*), was similarly enriched at the endogenous enhancer loci of all clusters (Fig. 2D,E). We infer that the distinct specificities did not stem from differences in initial enhancer strengths. The histone modifications H3K4me1 and H3K4me3 did however show differential and complementary trends: H3K4me1 was the highest enriched at cluster 5, followed by 4 and 3, whereas H3K4me3 was more highly enriched at enhancer clusters 1 and 2 (Figs. 2D,F,G). High H3K4me1 levels combined with high H3K27ac levels have been associated with distal regulatory regions or cell type-specific enhancers(*39*), suggesting that enhancers with the highest H3K4me1 levels (cluster 5) might be most specific to S2 cells. Analysing chromatin accessibility and H3K27ac levels in other cell types and tissues(*31, 40, 41*) (see methods) indeed revealed that enhancers from cluster 5 were highly cell type-specific (fig. S2). Interestingly, these enhancers were also the most strongly repressed by CoRest, CtBP and Rbf2 in our assays (Fig. 2B), suggesting that developmental or cell type-specific enhancers might intrinsically be more sensitive to repression by certain CoRs, while more globally active or housekeeping enhancers might be more resistant.

As active enhancers are known to function through a variety of different TFs, we hypothesised that their differential response to the CoRs might be linked to the specific TFs they bind. Using published TF ChIP-seq data, we indeed observed that prominent TFs were bound to enhancers from specific clusters and absent from others. The TFs DREF(*42*) and M1BP(*43*) for example bound almost exclusively to enhancers from clusters 1 and 2, with DREF preferring cluster 1 and M1BP cluster 2 (Figs. 2D,H,I). Trithorax-like(*44*) (Trl, also called GAGA factor or GAF) on the other hand was absent from these clusters and instead mainly bound to enhancers from cluster 3 and to a lower extent to cluster 4 (Figs. 2D,J). The distribution of these three TFs suggests that differential CoR sensitivities might relate to distinct TFs.

To identify associations between CoR sensitivities and TFs in a more comprehensive manner, we performed TF-motif enrichment analyses for the enhancers of the five clusters using the 6502 TF motifs from the iRegulon database(*45*). Consistent with the ChIP-seq results, motifs for DREF, M1BP and Trl were specifically enriched in clusters 1, 2 and 3, respectively (Fig. 2K). Additional motifs were also differentially enriched between the clusters: Rsc30 and E-box motifs were enriched in cluster 3, Sim motifs were enriched in cluster 4, while ETS and GATA motifs were enriched in cluster 5. Other prominent TF motifs were enriched over several clusters, including AP-1 (clusters 3-5), Ato (clusters 1,3,4) and CrebB (clusters 3,4). Taken together, the different enhancer clusters, defined by their sensitivity towards CoRs, associate with distinct chromatin features and TF motif content.

### Identification of TF motifs that predict sensitivity and resistance to each CoR

To directly test the association between TF motifs and sensitivity towards each of the CoRs, we evaluated the enrichment of TF motifs within enhancers that were sensitive (FDR < 0.05, logFC < -0.3) versus those that were resistant to each CoR (Fig. 3A, see methods). For CtBP, for example, we found that AP-1, Trl and GATA motifs were strongly enriched in sensitive enhancers, whereas DREF, Ohler1 and Mip120 motifs were enriched in the resistant enhancers (Fig. 3A; see fig. S3 for other CoRs).

**Fig. 3.**
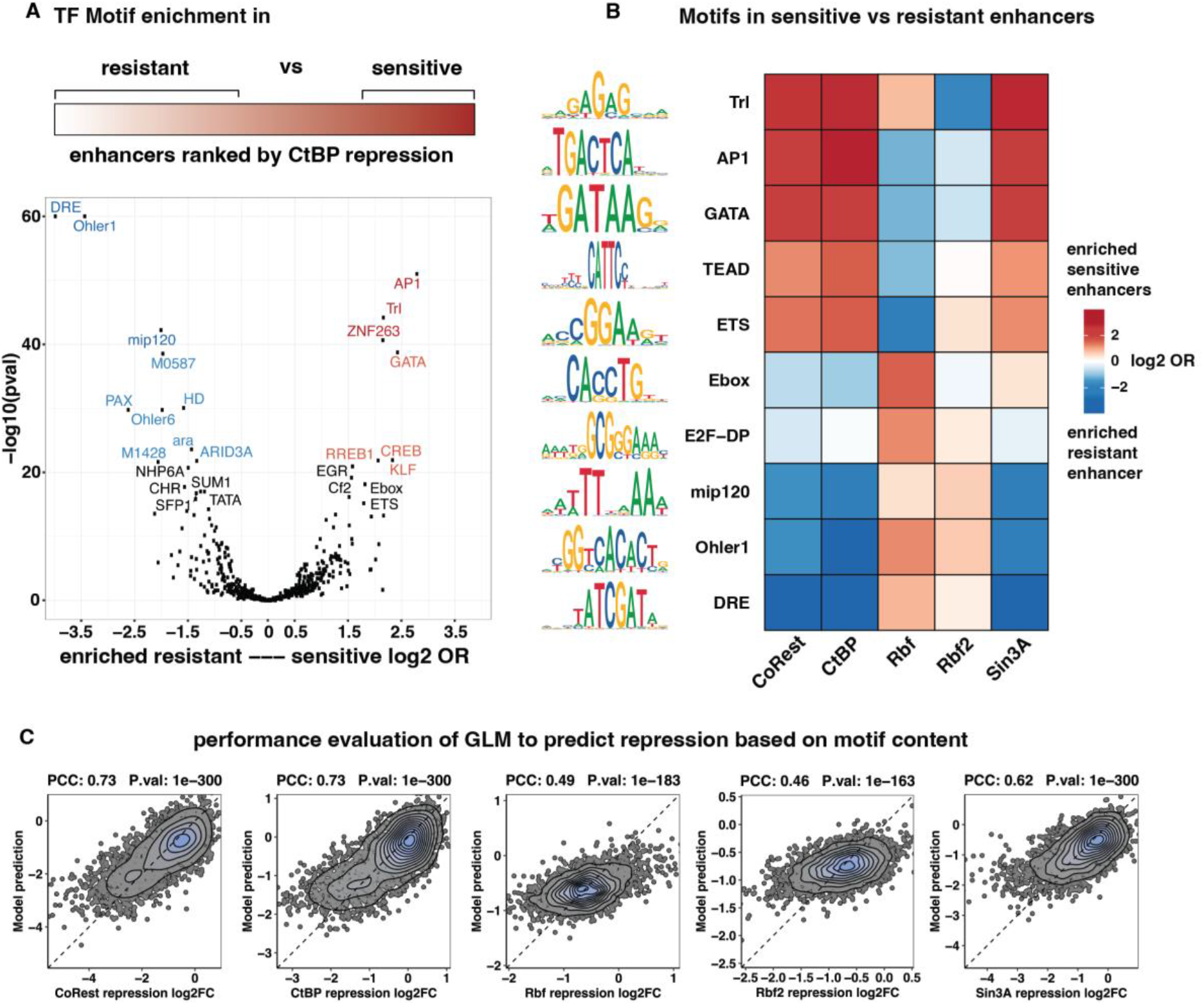
Identification of TF motifs that correlate with sensitivity or resistance to a given CoR. (**A**) Volcano plot of TF motifs enriched in enhancers that are either resistant to CtBP (left) or sensitive (right). Enrichment on x-axis log2(odds ratio counts sensitive/counts resistant), significance on y-axis –log10 Pval Wilcoxon Test. (**B**) Heatmap summarizing the enrichment of TF motifs in enhancers that are sensitive (red) or resistant (blue) to the given CoR. (**C**) Correlation plots of the measured repression by STARR-seq log2FC(Gal4-CoR/Gal4-GFP) (x-axis) and predicted repression by the GLM (y-axis) for each CoR. Pearson correlation coefficient and significance for each model are displayed on top.

Motif enrichment profiles across all five CoRs revealed an intricate relationship between enhancer-CoR sensitivity and TF motifs with highly distinctive enrichments (Fig. 3B). Consistent with the conserved role of Rb proteins as part of the DREAM complex(*27, 46*), E2F, DP, and Mip120 motifs were enriched in Rbf sensitive enhancers. Interestingly, these three motifs as well as DREF and M1BP motifs were enriched in enhancers that were resistant to CoRest, CtBP and Sin3A repression (Fig. 3B). On the other hand, motifs for the developmental TFs GATA, AP1 and TEAD were enriched in enhancers sensitive to CoRest, CtBP and Sin3A and resistant to Rbf and Rbf2 (Fig. 3B). Furthermore, Trl and ETS motifs were specifically enriched in enhancers that were resistant to Rbf2 and Rbf respectively, while sensitive to all other CoRs (Fig. 3B). These results indicate that, for each CoR, certain TF motifs are specifically and strongly enriched in sensitive and resistant enhancers.

Given the differential motif enrichments, we next tested whether TF motif content is predictive of an enhancer’s CoR sensitivity. For this, we trained a Generalized Linear Model (GLM)(*47*) using TF motif counts as features and enhancer sensitivity to a given CoR as response variable (see Methods). For each CoR, using 10-fold cross-validation, the models based on motif counts performed well and were able to predict an enhancer’s sensitivity and resistance to a given CoR (PCCs between 0.46 and 0.73, Pvals < 2.2*10^−16^; Fig. 3C). Overall, these results establish a strong association between CoR sensitivity and TF motif content that is predictive and might correspond to causal relationships.

### Distinct transcription factor motifs are required for resistance against specific repressors

We noticed that DREF TF motifs were enriched in enhancers that were resistant to CoRest and CtBP (Fig. 2K, 3B). Indeed, enhancers bound by DREF, such as enhancers near the *RYBP* and *jupiter* genes (Fig. 4A), are specifically resistant to repression by CoRest and CtBP but sensitive to repression by Rbf (Fig. 4A). Ranking all enhancers based on their sensitivity to repression by CoRest or CtBP confirmed that resistant enhancers were significantly (Pval < 2.2*10^−16^) enriched for DREF motifs and bound by DREF according to ChIP-seq(*42*) (Fig. 4B). Given the correlation between CoRest and CtBP resistance and presence of DREF, we considered that DREF might protect enhancers against CoRest and CtBP-mediated repression. To test whether DREF motifs are indeed required for the resistance, we selected 65 DREF-motif containing enhancers, mutated the DREF motifs and assessed the enhancers’ sensitivity to repression (see Methods). The mutated enhancers were significantly more sensitive to repression by CoRest and CtBP (Pvals = 4.1*10^−11^ & 2.5*10^−10^), while their sensitivity towards Rbf did not change (Figs. 4C,F). We infer that DREF motifs are required for resistance to CoRest and CtBP.

**Fig. 4.**
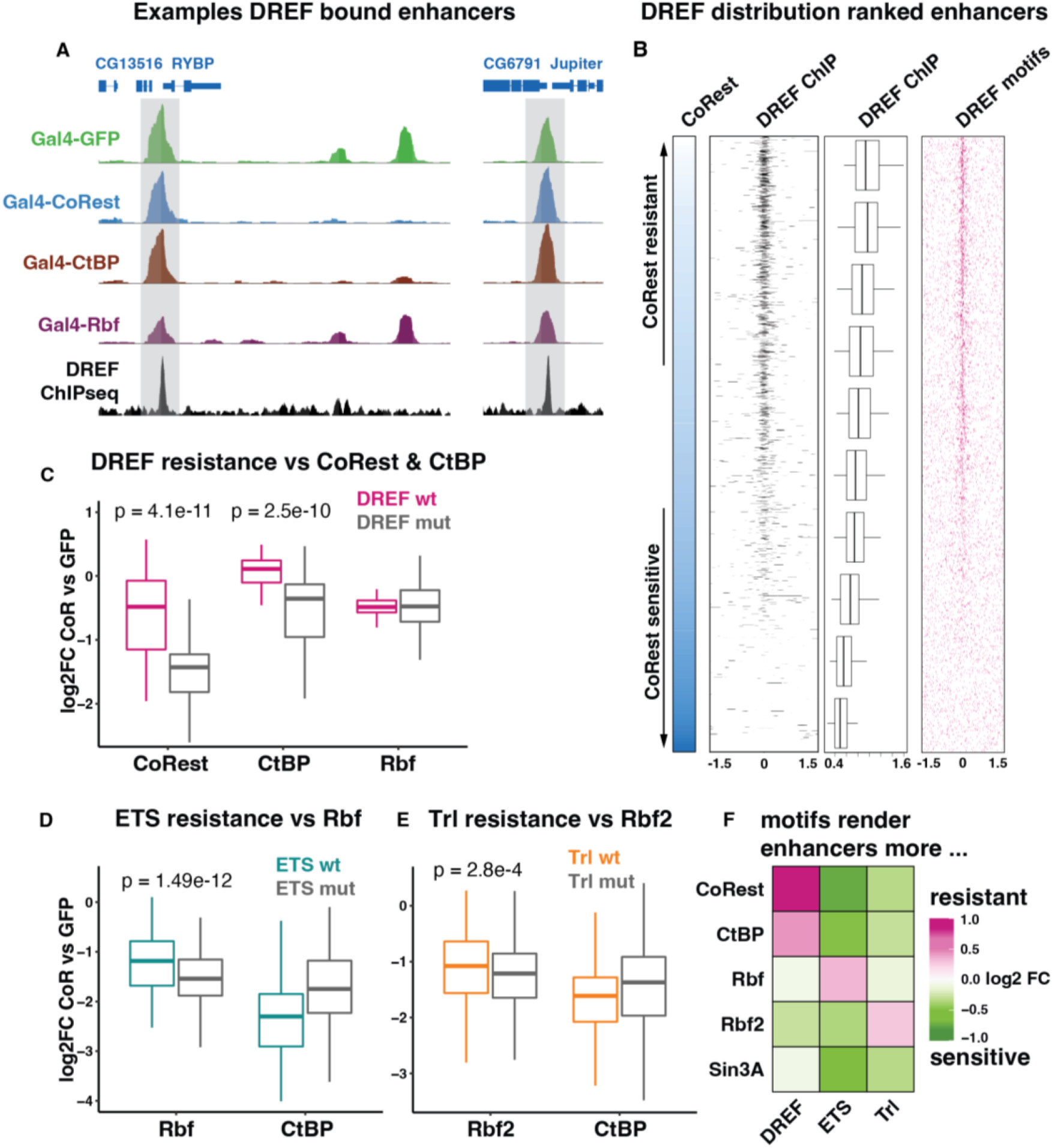
Testing requirement of specific TF motifs to confer resistance against distinct repressors. (**A**) UCSC screenshot of normalized STARR-seq tracks displaying DREF bound and unbound enhancers around the *RYBP* and *Jupiter* genes. Top to bottom; STARR-seq with Gal4-GFP, Gal4-CoRest, Gal4-CtBP, Gal4-Rbf and ChIP-seq against DREF. (**B**) Heatmap of all enhancers ranked based on their sensitivity to CoRest. ChIP-seq signal of DREF is displayed, quantified in boxplots (Pval < 2.2*10^−16^) and the occurrence of DREF motifs is plotted. (**C**) Boxplots showing the log2FC of enhancer activity when CoRest (left), CtBP (mid) or Rbf (right) is recruited over GFP as control. Each enhancer is present as original (pink) and with its DREF motifs mutated (grey), n = 65. (**D**) Boxplots showing the log2FC of enhancer activity when Rbf (left) or CtBP (right) is recruited over GFP as control. Each enhancer is present as original (cyan) and with its ETS motifs mutated (grey), n = 157. (**E**) Boxplots showing the log2FC of enhancer activity when Rbf2 (left) or CtBP (right) is recruited over GFP as control. Each enhancer is present as original (orange) and with its Trl motifs mutated (grey), n = 127. (**F**) Heatmap summarizing the effects of motif mutations on enhancer sensitivity to the different CoRs (log2FC of repression wild type enhancers over repression when specific motif is mutated).

Several other TF motifs were predicted to confer resistance to repression by specific CoRs. ETS-family motifs, for example, were specifically enriched in enhancers that were resistant to Rbf repression (Fig. 3B). We mutated the ETS motifs in 157 enhancers and indeed observed increased sensitivity towards Rbf-mediated repression (Pval = 1.49*10^−12^, Fig. 4D). Remarkably, the opposite trend was observed for the other CoRs, which were all able to repress the wildtype enhancers containing ETS motifs better than their mutated counterparts, suggesting that ETS TFs are in general sensitive to repression (Figs. 4D,F). Similarly, Trl motifs were enriched in enhancers that were specifically resistant to Rbf2 repression (fig. S4) and mutating these motifs in 127 enhancers led to a slight but specific increase in sensitivity towards Rbf2, but not towards the other CoRs (Fig. 4E,F).

In general, specific TF motifs are required to protect resistant enhancers against a given CoR. As these motifs and enhancers are however still sensitive to other CoRs, an intricate pattern emerges (Figs. 4F & S4-5), which suggests that the interplay between repressors and activators can be highly specific.

### Specific transcription factor motifs can turn sensitive enhancers into resistant ones

Given that specific TF motifs are required for CoR resistance, we wondered whether these motifs are also sufficient to cause CoR resistance. To test this, we introduced two “resistant” TF motifs (10 bp each) into CoR-sensitive enhancers at positions deemed unimportant for enhancer activity(*48*) (see methods and S.table x) and evaluated the enhancers’ sensitivity to repression (Fig. 5A). As a control for the addition of motifs, we also introduced two neutral control TF motifs at the same positions (of the ACE2 and FOX TFs), which were predicted to not impact the sensitivity to repression (see methods).

**Fig. 5.**
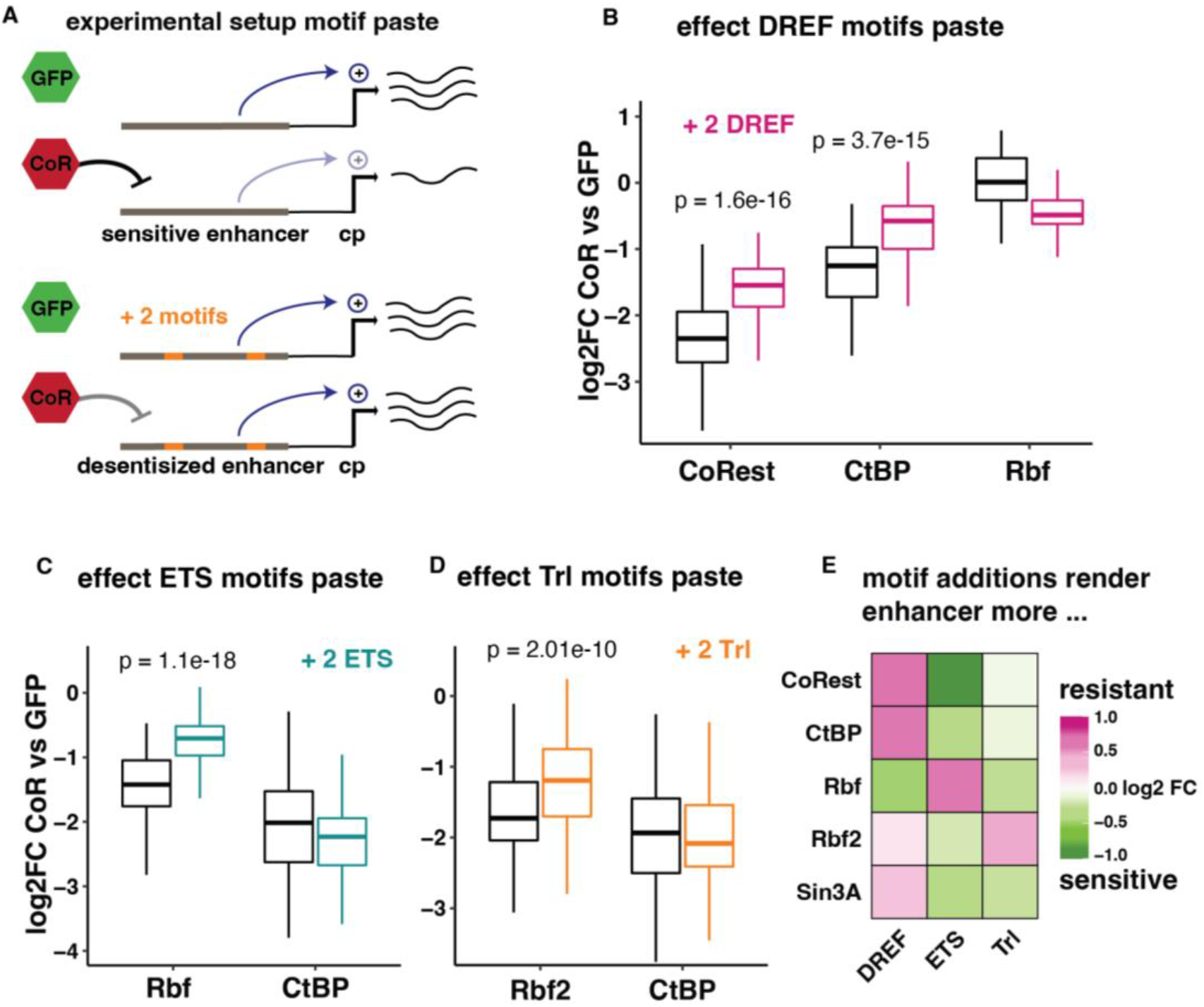
Effects of TF motif addition on enhancers sensitivity to repression. (**A**) Experimental setup of the motif paste experiments. Two TF motifs that were predicted to add resistance are pasted into the sequence of sensitive enhancers (see methods). The activity of original and motif paste enhancers is measured by UAS-STARR-seq under control (Gal4-GFP) and repressor (Gal4-CoR) conditions (see Methods and Data S2). (**B**) Boxplots showing the log2FC of enhancer activity when CoRest (left), CtBP (mid) or Rbf (right) is recruited over GFP as control. Each enhancer is present as original (black) and with two DREF motifs are pasted (pink), n = 110. (**C**) Boxplots showing the log2FC of enhancer activity when Rbf (left) or CtBP (right) is recruited over GFP as control. Each enhancer is present as original (black) and with two ETS motifs are pasted (cyan), n = 107. (**D**) Boxplots showing the log2FC of enhancer activity when Rbf2 (left) or CtBP (right) is recruited over GFP as control. Each enhancer is present as original (black) and with the addition of two Trl motifs (orange), n = 93. (**E**) Heatmap summarizing the effects of the motif additions on enhancer sensitivity to the different CoRs (log2FC of repression with motif addition vs repression without motif addition).

Since DREF motifs were required for resistance to CoRest and CtBP-mediated repression (Fig. 4C), we tested whether they were sufficient to render CoRest and CtBP-sensitive enhancers resistant. Overall, the introduction of two DREF motifs was sufficient to protect the enhancers from CoRest and CtBP-mediated silencing (Pval = 1.59*10^−16^ & 3.71*10^−15^, n=110, Fig 5B) and this protection was specific and due to the DREF motifs, as control motifs had no effect on sensitivity (fig, S5A). Similarly, ETS motifs were necessary (Fig. 4D) and sufficient (Fig. 5C) to desensitize enhancers from repression by Rbf (Pval = 1.06*10^−18^, n=107), and Trl motifs were necessary (Fig. 4E) and sufficient (Fig. 5D) to specifically generate Rbf2-resistant enhancers (Pval = 2.01*10^−10^, n=93).

In each case, the gain in resistance due to the addition of selected TF motifs was highly CoR specific, as these new motifs had little effect on the repression by other CoRs or could even sensitize the enhancers to other CoRs (Figs. 5E & S6). The addition of ETS or Trl motifs for example made enhancers more sensitive to all other CoRs except for Rbf or Rbf2 respectively (Figs. 5E & S5), while DREF motifs increased sensitivity towards Rbf (Pval = 6.36*10^−14^). Hence, introducing specific TF motifs changed the enhancer’s sensitivity to repression by a given CoR in a specific and predictable manner.

Taken together, we conclude that specific TFs are necessary and sufficient to confer resistance to repression by a given CoR, while other CoRs are able to repress these TFs, indicating that certain TFs can directly counteract some CoRs and their repressive function or that different modes of transcriptional activation require different modes of repression.

## Discussion

In this work, we show that transcriptional enhancers are differentially susceptible to active repression by prominent CoRs. We mapped the regulatory specificities between active enhancers and five CoRs on a genome-wide scale and discovered that enhancers across the genome are repressed by just a subset of specific CoRs. This additional layer of regulatory specificities enables the differential repression of closely spaced enhancers and genes as they frequently occur in the *Drosophila* genome. At the *fs(1)h/mys* locus for example, CoRest could repress the intronic *mys* enhancer without affecting the neighboring *fs(1)h* gene, whereas Rbf could do the opposite (Fig. 6A left). Similarly, at the *kay/fig* locus some CoRs like CtBP could repress both closely spaced enhancers, whereas Rbf2 could specifically repress the enhancer closest to *fig* but not the others (Fig. 6A right).

**Fig. 6.**
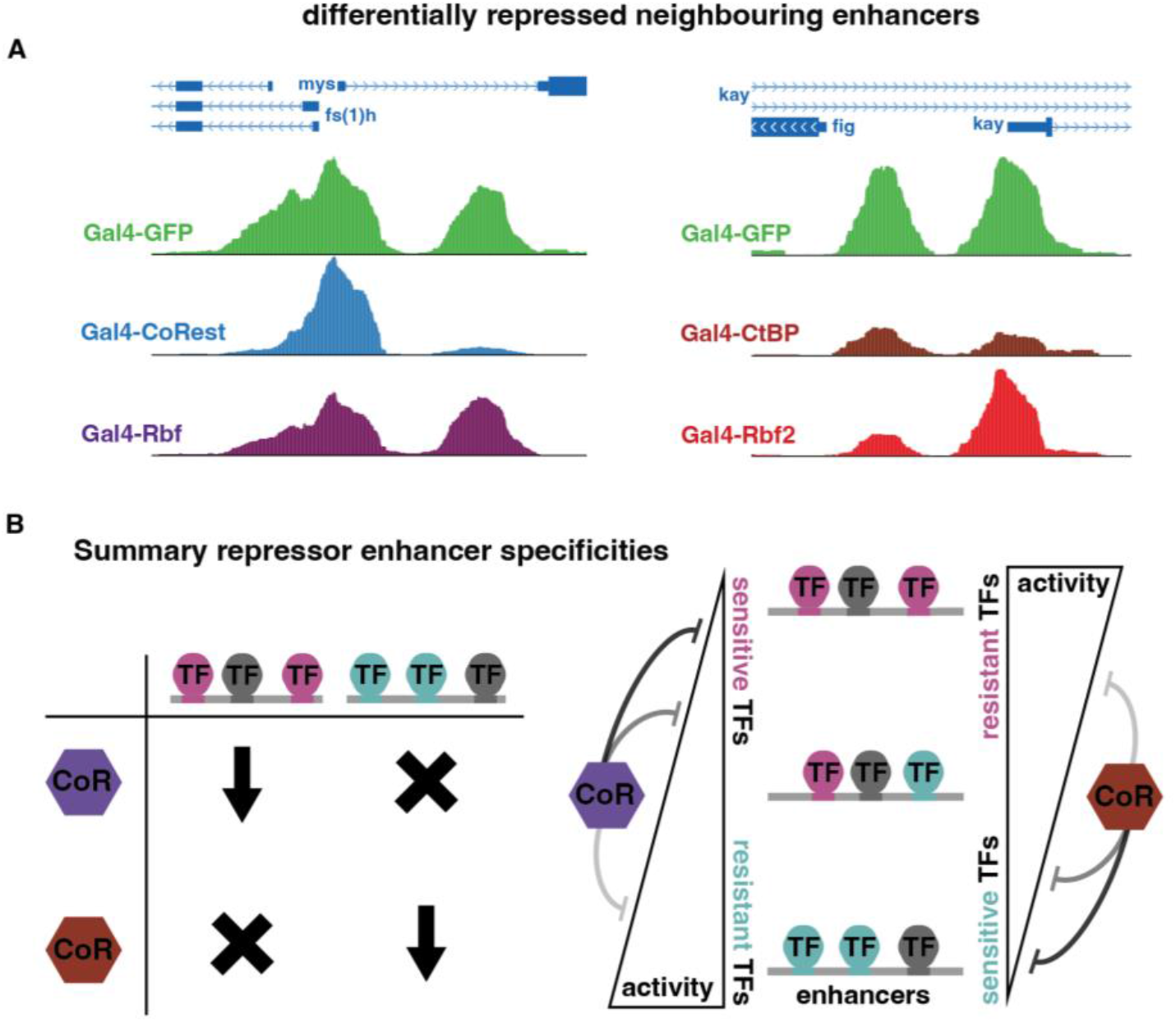
Implications of repressor-activator specificity. (**A**) Example regions with two closely spaced but differentially repressed regulatory regions. Left enhancers differentially repressed by CoRest and Rbf near the *fs(1)h* and *mys* genes and right enhancers differentially repressed by CtBP and Rbf2 near the *fig* and *kay* genes. (**B**) Summary schematic of uncovered specificities between repressors and activators. (**C**) Summary schematic of the additional layer in transcriptional regulation. Depending on motif content (pink, cyan or grey) enhancers can be sensitive to one CoR while resistant to another one and different combinations lead to intermediate repression.

The uncovered specificities between repressors and enhancers not only suggest that repression occurs via distinct mechanisms, but also reveal a previously unappreciated layer of “resistance” against repression (Fig. 6B). Enhancers that are sensitive or resistant towards a given CoR display clear differences in their TF motif content, and motif mutagenesis experiments changed enhancer sensitivities in a predictable manner. Specific motif combinations of activating and repressive TFs, together with the regulatory proteins present in each cell type, will largely determine when and where a regulatory region is active or repressed(*6, 49*). In addition, the ability of TFs to confer resistance also allows for enhancers and genes to be activated by de-repression via TF recruitment, as we demonstrate by motif addition (Fig. 5). This additional layer of regulation circumvents the requirement to remove the involved repressors and the interplay between TFs and CoRs provides more flexibility to the regulatory system (Fig. 6C).

Importantly, the fact that one mode of activation can be sensitive to one repressor but resistant to another implies that distinct activation and repression mechanisms must converge. Active repression might directly interfere with specific factors or defined steps of transcriptional activation. Certain CoRs might for example inactivate specific TFs but not others (e.g. via posttranslational modifications(*50, 51*)), or counteract the TFs’ downstream activating mechanisms. Alternatively, certain TFs might directly counteract a repressor’s function or bypass the rate-limiting step of initiation- or elongation controlled by the CoR. Discerning the distinct mechanisms of activation and repression and how they intersect and coordinate will be of great future interest.

## Acknowledgments

We thank the core facilities at the VBCF for their excellent support, in particular the molecular biology service, NGS facility and HPC team. We thank Bernardo P. de Almeida for assisting with the oligonucleotide library design, A. Andersen (Life Science Editors) for comments on the manuscript and G. Hulselmans and S. Aerts (KU Leuven) for sharing the TF motif PWM collection.

## Author contributions

Conceptualization: JJ, AS

Methodology: JJ, MP, CW

Investigation: JJ, AS

Visualization: JJ

Funding acquisition: JJ, AS

Supervision: AS, JJ

Writing – original draft: JJ, AS

Writing – review & editing: JJ, AS

## Data and materials availability

All raw and processed NGS data related to this paper has been deposited to GEO: GSE215143. The UAS-STARR-seq tracks can be viewed on the UCSC browser: https://genome.ucsc.edu/s/jelle.jacobs/rep_STARR_gw_norm_merged

## Supplementary Materials

Materials and Methods

Figs. S1 to S6

