## Supplemental Figs and M&M for "Widespread regulatory specificities between transcriptional corepressors and enhancers in *Drosophila*"

**This PDF file includes:**

Materials and Methods
Figs. S1 to S6

### Materials and Methods

#### UAS-STARR-seq library cloning

In original STARR-seq(*1*), a library of genomic DNA fragments is cloned downstream of an open reading frame (ORF) and a core promoter. Fragments that harbour enhancer activity will stimulate transcription from the core promoter and generate transcripts that contain the enhancer sequence. STARR-seq transcripts are captured, sequenced and mapped back to the genome, allowing for the simultaneous discovery of enhancers and their relative strength. In order to recruit selected CoRs to this enhancer library, a 4xUAS sequence (cggagtactgtcctccgagcggagtactgtcctccgaggccgccgcccggagtactgtcctccgagcggagtactgtcctccgag) was cloned into the original *Drosophila* STARR-seq vector(*1*) so that the 4xUAS sequence would be 17bp downstream of the enhancer library allowing for the recruitment of Gal4-CoR near then enhancers; UAS-STARR-seq. All libraries were generated with both the *Drosophila* synthetic core promoter (DSCP) and the Rps12 core promoter(*2*).

For the *Drosophila Melanogaster* genome-wide (gw) library, sheared genomic DNA fragments were processed as previously described(*1*) and cloned into the UAS-STARR-seq vectors using In-Fusion HD cloning reactions (In-Fusion HD Cloning Kit, Clonetech; Cat No. 639650).

For the oligonucleotide designed library (olig) (used in Figs. 4C-F & 5) 249bp designed enhancer fragments (both wt and motif mutants) were ordered at Twist Bioscience, PCR amplified and cloned into the UAS-STARR-seq vectors using Gibson cloning (New England BioLabs, catalog no. E2611S).

Libraries were transformed in electrocompetent MegaX DH10B-T1 cells and grown in 12L (gw) or 6L (olig) Luria-Bertani medium with 100ug/ml ampicillin. After growing ~12h libraries were purified with a Qiagen Plasmid Plus Giga Kit (catalog no. 12991).

#### Spike-in controls

Spike-ins were created similar to the UAS-STARR-seq libraries above, with the important difference at this time the original *Drosophila* STARR-seq vector(*1*) was used (so without 4xUAS site). As library sheared pieces of the *Drosophila Pseudoobscura* genome were used; Bacterial Artificial Chromosomes 1, 2, 3, 4, 7, 8, 18 & 19). The activity of 18 *D. Pseudoobscura* enhancers residing in these BACs (see sup table spike-in reads) was used to normalize the UAS-STARR-seq libraries.

#### Transfection of UAS-STARR-seq

The gw UAS-STARR-seq libraries with DSCP and Rps12 core promoters were premixed in equimolar ratios and 1% *D. Pseudoobscura* spike-in was added. The olig libraries with DSCP and Rps12 core promoters were kept separate and 1% of *D. Pseudoobscura* spike-in was added. Per OC-400 (electroporation unit), 20ug of the UAS-STARR-seq library was mixed with 20ug of a vector that expressed either Gal4-GFP or one of the Gal4-CoRs in S2 cells (expression vector pAGW-GAL4-DBD with the ORF of GFP, CoRest, CtBP, Rbf, Rbf2 and Sin3A as in ref(*3*)).

*Drosophila* S2 cells were cultured under standard conditions(*1*) and harvested while in the growth phase. Per electroporation unit (OC-400) 200 x $10^6$ cells were electroporated with 40ug of the UAS-STARR-seq/spike-in/Gal4-CoR mix, using the MaxCyte-STX system at a density of 50 x

$10^6$ cells per 100 µl. For each individual sample of the gw screens, 2 OC-400 were performed, and cells were afterwards processed together, while the olig screens only required 1 OC-400 per sample. The rest of the UAS-STARR-seq protocol is the same as the regular STARR-seq protocol(*1*).

**Illumina sequencing and mapping**

The VBCF NGS facility performed the next-generation sequencing on a NextSeq 550 (gw screens, paired end 36bp) or NovaSeq SP (olig screens, PE 150bp) platform, following the manufacturer's protocol, using standard Illumina i5 and i7 indexes.
Processed UAS-STARR-seq reads from gw screens were mapped to the *Drosophila Melanogaster* (dm3) genome and subsequently to the artificial *Drosophila Pseudoobscura* BAC genome (spike-in), using Bowtie v.1.2.2.(*4*). PE 36bp reads with a maximal insert size of 2 kb and up to three mismatches were kept.
The olig screens were mapped to a synthetic reference genome containing the 249bp sequences with both wildtype and mutant enhancers, only exact matches were kept. Subsequently the reads were mapped to the artificial *Drosophila Pseudoobscura* BAC genome like in the gw screens (spike-in).

**Differential enhancer activity analysis**

Reads that mapped to the 18 *Drosophila Pseudoobscura* BACs spike-in enhancers were quantified (see sup table spike-in reads). As we always used the same amount of spike-in and the spike-ins were not affected by the expressed CoRs (no 4xUAS sequence) they severed as internal controls to normalize the samples to. Practically, first a spike-in reference was generated by taking the average spike-in counts per enhancer across all samples. Then a linear regression coefficient between this reference and each sample was calculated and used as normalization factor for the UAS-STARR-seq libraries.
For the gw screens, active enhancers were identified with macs2(*5*) callpeak using –nomodel and –fe-cutoff 4 as additional parameters. Enhancers were called on all 12 samples separately, and their genomic coordinates were merged into one enhancer backbone file. STARR-seq reads falling into these enhancers were quantified for every sample using countOverlaps from the GenomicRanges(*6*) R-package, obtaining a raw counts matrix. The raw counts matrix was then normalized using the spike-in normalization factors and the normalized matrix was used to perform differential enhancer activity analysis. To be able to quantify repression, enhancer with at least 200 normalized reads in both control (Gal4-GFP) samples were selected (3094 enhancers, see normalized counts table). The differential expression package EdgeR(*7*) was used to quantify the differences in enhancer activity between control (Gal4-GFP) and the different Gal4-CoRs, each time using both biological replicates.
For the oligo screens read were mapped to every oligo, directly generating the raw counts matrix. Further processing was done in the same way as the genome-wide screens (see olig count table).

**Luciferase reporter assay**

The pGL3 Luciferase Reporter Vector (Promega) was used to clone the validation vectors. The promoter was replaced by the DSCP or Rps12 promoters as in(*2*) and the 4xUAS site followed by

the tested enhancer was cloned upstream of the promoters. Following enhancer were cloned (dm3); srp-DSCP chr3R:11810070-11810909, CG2116-Rps12 chrX:8023445-8024243 and kay/fig-DSCP chr3R:25606440-25607231.
Luciferase assays were performed as previously described(*1*) with the following transfection mix: 100ng luciferase plasmid, 10ng Gal4-CoR expression plasmid and 5ng Renilla plasmid per well. Each reaction was done in triplicate (technical replicates) and performed in at least two biological replicates.

**Clustering of enhancers using SOTA**

The enhancers from the genome-wide screens were clustered based on their differences in sensitivity towards the 5 CoRs log2(enhancer activity Gal4-CoR/ enhancer activity Gal4-GFP, as calculated above). Using a self-organizing tree algorithm (SOTA)(*8*) with the Pearson correlation coefficient as distance metric, we grouped the enhancers into five clusters, explaining 54% of the total variance. The sensitivity of enhancers towards the CoRs in each cluster was summarized in boxplots displaying the log2FC in enhancer activity over GFP. Boxplots display the median, hinges correspond first and third quartiles and whiskers to the largest value maximally 1.5* the inter-quartile-range.

**Public datasets**

Following previously published datasets were reanalyzed in this study: ChIP-seq for the chromatin marks H3K27ac(*9*), H3K4me1(*10*) and H3K4me3(*11*) in S2 cells and H3K27ac in D. Melanogaster 3$^{rd}$ instar larval eye-antennal discs(*12*), ChIP-seq for TFs DREF(*13*) in Kc167, M1BP(*14*) and Trl(*15*) in S2 cells. Accessible chromatin by ATAC-seq in S2(*16*), in D. Melanogaster 3$^{rd}$ instar larval brains and eye-antennal discs(*17*) and by DHS in OSCs(*1*). Raw sequencing data was re-mapped to the dm3 genome assembly using Bowtie v.1.2.2.(*4*).

**Chromatin marks and TF ChIP enrichment over clusters**

Heat maps that visualize the coverage of genomic data were centered on the STARR-seq enhancers and the custom R script from(*18*) was used to plot the data. For the quantification of ChIP-seq and accessible chromatin data we used for every dataset the ChIP-peak signal inside each enhancer and over 1000 negative control regions. We then grouped the enhancers based on the repression clusters or repressor ranking and visualized the data in standard boxplots, showing the median, hinges correspond first and third quartiles and whiskers to the largest value maximally 1.5* the inter-quartile-range.

**TF-motif enrichment analyses**

The DNA sequence of each enhancer was scored for the presence of TF motifs using the Bioconductor package motifmatchr(*19*) counting each motif that passed the Pval cutoff of $1e^{-4}$. In total 6502 motif position weight matrices (PWMs) from a collection of databases described in ref(*20*) were used to score each enhancer. Motifs that were significantly enriched in one enhancer cluster compared to the other clusters were identified using the Fisher's exact test. The log2 odds ratio of significant motifs was used to plot their enrichment or depletion over the enhancer clusters

(Fig. 2K). In Fig.3 for each CoR individually, we compared all sensitive enhancers (log2FC < -0.3 & FDR < 0.05) to that CoR to all resistant enhancers (log2FC > -0.3). Significantly enriched motifs in either sensitive or resistant enhancers were identified using the two-sided Wilcoxon rank-sum test and the log2 odds ratio was calculated for each motif.

**Generalized linear model to predict enhancer sensitivity to CoRs**

To evaluate how well the enhancers motif content can predict its sensitivity towards a given CoR, a generalized linear model was fitted to the data. Motif counts were used as input matrix and sensitivity of the enhancers towards each CoR log2(enhancer activity CoR/GFP) was the response vector. Practically, the R package glmnet(*21*) was used with $\alpha = 1$ to train and evaluate the models using 10 fold cross validation.

**Designed oligonucleotide (TWIST) library with motif mutants and motif paste experiments**

For the motif mutant and paste experiments, we computationally designed a 300-mer oligonucleotide library as was done in(*22*), that was synthesized by Twist Bioscience. For the motif mutant experiments, we selected wild-type enhancers containing DREF (ATCGAT) n=189, ETS (CCGGAA) n=173 and Trl (GAGAG) n=674 motifs. We then mutated these motifs in the enhancers to GCTTAA, TTACGC and AAGTA respectively, sequences that were predicted to not have regulatory activity in S2 cells.
For the motif paste experiments, we selected 20 wild-type enhancers that were sensitive to one or multiple CoRs and pasted two "resistant" (DREF = CTATCGATAG, ETS = ACCGGAAGTG or Trl = AGAGAGAGAG) or control (FOX = GTAAACAAAC, ACE2 = ATGCTGGTTC) 10bp TF motifs into regions that were predicted to be unimportant for enhancer activity using DeepSTARR(*22*) (see Table Motif_paste_experiments with coordinates for each enhancer and where the motifs were pasted inside the enhancers). To further evaluate the direct effects that each pasted TF motif has on repression, we increased the number of enhancer pairs to compare (no "resistant" motifs vs "resistant" motifs pasted) by changing the enhancers backbones. Basically, for every initial enhancer, DeepSTARR(*22*) selected the regions that are important for enhancer function (containing different activator motifs) and those regions were copied into 10 neutral backbones (see Table Motif_paste_experiments). In these newly generated enhancers, we then pasted the "resistant" and control motifs in the same locations as done for the initial enhancers. The exact sequence of each oligo can be found in the supplementary files of the GEO submission (GSE215143_olig_repSTARR_seq).
As mentioned above in STARR-seq library cloning, we cloned two libraries, one with the DSCP and one with the Rps12 core promoters and – in contrast to the screen with the genome-wide libraries above – screened them separately, two independent replicates per core promoter.

**Analysis of motif mutant and paste experiments**

For the downstream analysis, we required that each individual oligo in the UAS-STARR-seq library was > 20 times present in the input and that the enhancer activity under control (Gal4-GFP) conditions was > 20 normalized reads in both replicates. For each comparison between wild-type and mutant enhancer, we required that both made these cutoffs. These stringent filters allowed us to confidently evaluate repression and the direct effects of motif mutants and additions.

As DREF motif-containing enhancers predominantly activated the Rps12 (hk) CP, while ETS motif-containing enhancers predominantly activated the DSCP (dev) CP, we evaluated their effects using the respective CP. Trl-motif-containing enhancers activated both CPs to similar extents, hence we merged the libraries and assessed CoR-mediated repression for the merge dataset. Differential enhancer activities were calculated as described above for the genome-wide libraries.

For the Trl mutant analysis, we selected all enhancers that on top of Trl motifs had also endogenous Trl binding (normalized ChIP-seq(*15*) signal above 8) retaining 127 high confident Trl bound enhancers and their mutant counterparts. For ETS (n=157) and DREF (n=65) all enhancers where both wt and mutant passed the cutoffs were retained. For the motif paste experiments we had 110 enhancer pairs with DREF motifs pasted, 107 with ETS motifs and 93 with Trl motifs that passed the cutoffs.

The effects on enhancer sensitivity (log2FC CoR/GFP) are visualized in boxplots, showing the median, hinges correspond first and third quartiles and whiskers to the largest value maximally 1.5* the inter-quartile-range. Significant differences between wt and motif mutants or additions were calculated using the paired Wilcoxon rank-sum test. The effect that a given motif has on the sensitivity of enhancers towards all tested CoRs is summarized in final heatmaps (Fig. 4F&5E). The change in sensitivity is calculated as the log2 of the average repression of enhancer with motif (CoR/GFP) over the average repression of the same enhancers without motif (CoR/GFP). Positive log2FC (pink) indicate the selected motif renders the enhancers more resistant against repression by that CoR, while negative values (green) indicate it renders the enhancers more sensitive to repression.

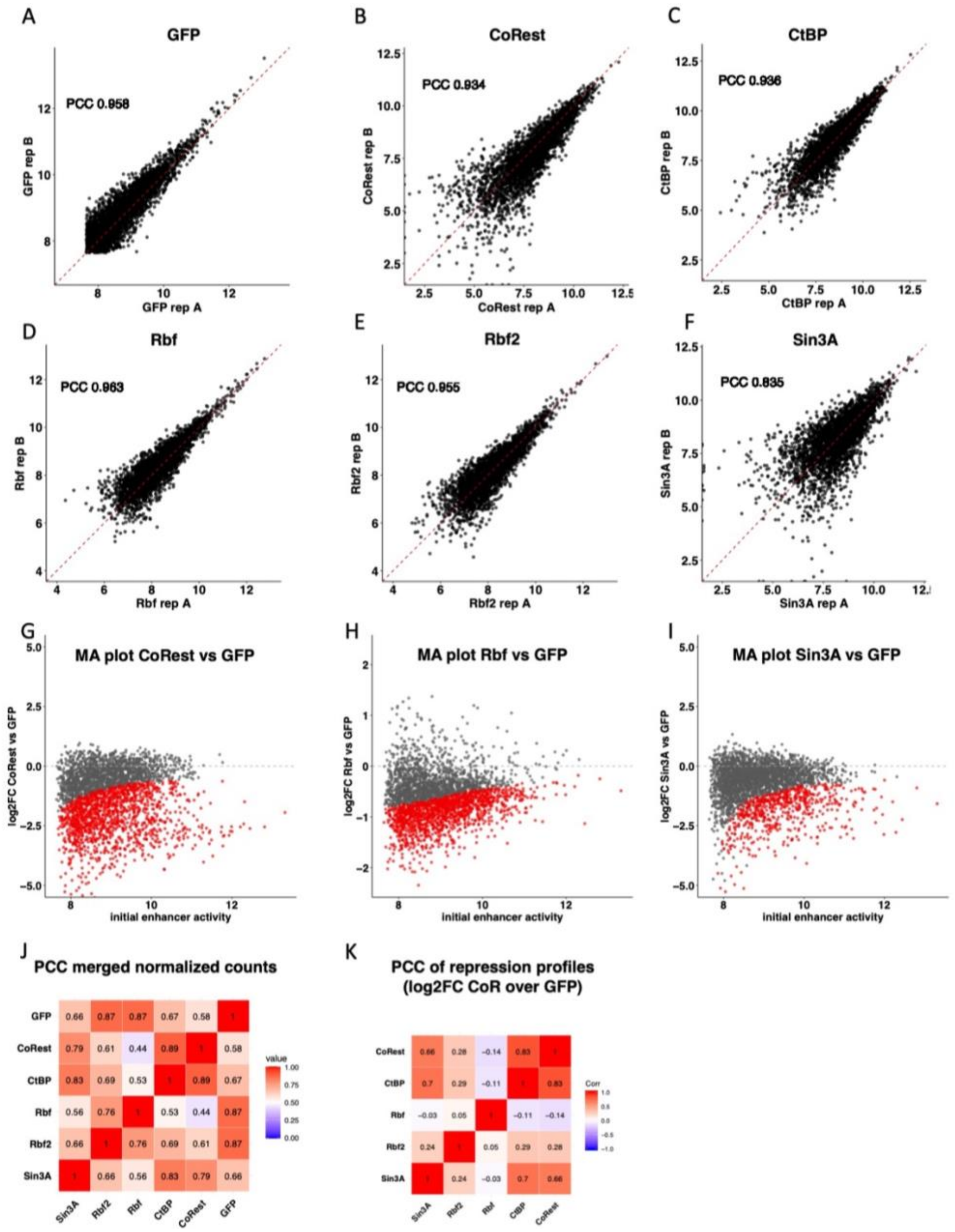


**Fig. S1. Correlation plots between biological replicates and PCC between the different repressors** (**A-F**) Correlation plots of the 3094 enhancers, each time two biological replicates are shown for when Gal4-GFP is recruited and the Gal4-CoRs; CoRest, CtBP, Rbf, Rbf2 and Sin3A. (**G-I**) MA-plots visualizing the repression of the three remaining CoRs; CoRest, Rbf and Sin3A. Initial enhancer activity (log2 normalized reads under Gal4-GFP) on x-axis and repression by CoR log2FC(enhancer activity Gal4-CtBP vs Gal4-GFP) on y-axis, n=3094. (**J**) Heatmap displaying the Pearson Correlation Coefficient between enhancer activity (merged spike-in normalized counts, 3094 enhancers) of all Gal4-CoR and Gal4 GFP screens. (**K**) Heatmap displaying the Pearson Correlation Coefficient between enhancer repression by the 5 CoRs (log2FC enhancer activity CoR / enhancer activity GFP).

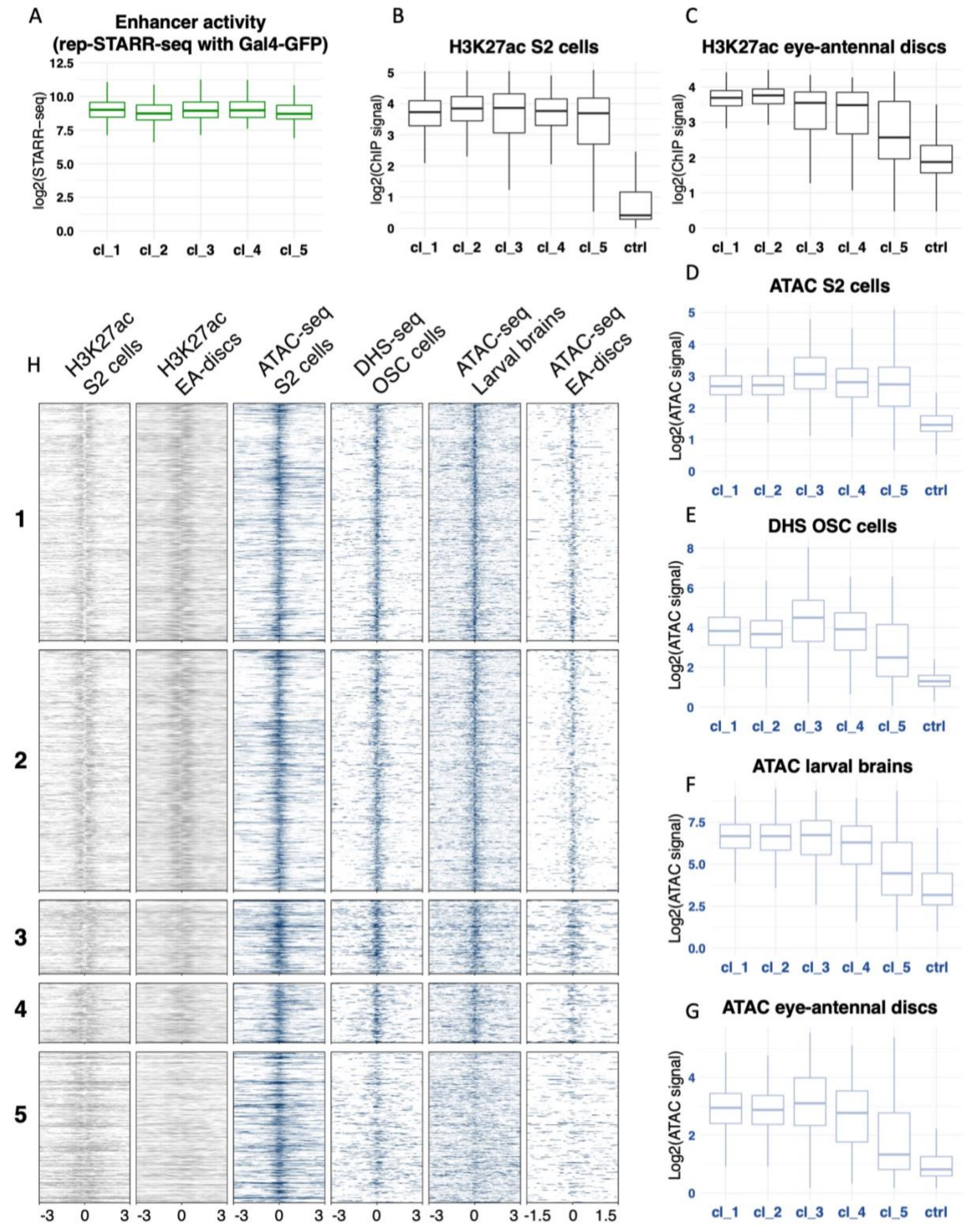


**Fig. S2. Distribution of endogenous chromatin marks and chromatin accessibility over the 5 repression clusters.** (**A**) Boxplot of enhancer activity under control (Gal4-GFP) condition. (**B-G**) Boxplots summarizing ChIP-seq or open chromatin signals over the different enhancer clusters, including negative control regions (1000 regions in the genome with no enhancer activity). (B)

H3K27ac in S2 cells, same as in main Fig 2 E. (C) H3K27ac in Drosophila 3$^{rd}$ instar eye-antennal discs. (D) ATAC in S2 cells. (E) DHS in Ovarian Somatic Cells. (F) ATAC in Drosophila 3$^{rd}$ instar larvae brains (G) ATAC in Drosophila 3$^{rd}$ instar eye-antennal discs. (H) Heatmap visualizing the ChIP- and open chromatin signals over the 5 enhancer clusters.

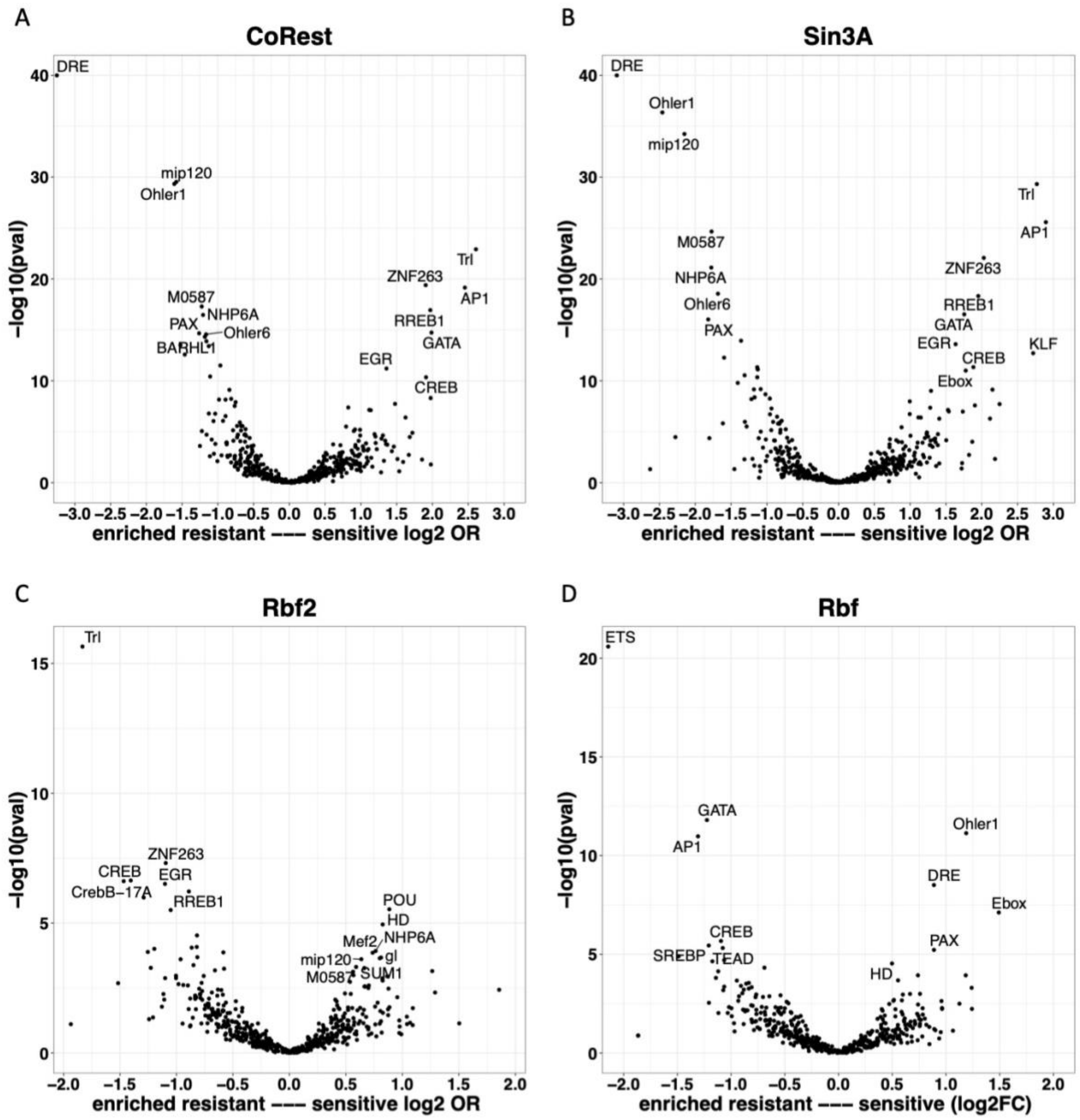


**Fig. S3. TF motif enrichment in resistant and sensitive enhancers to each CoR**

(**A-D**) Volcano plots of TF motifs enriched either in enhancers that are resistant to the given CoR (left) or enriched in sensitive enhancers (right). Enrichment on x-axis log2(odds ratio motif in sensitive enhancers vs in resistant), significance on y-axis –log10 P.val Wilcoxon Test. (A) For Gal4-CoRest, (B) Gal4-Sin3A, (C) Gal4-Rbf2 and (D) Gal4-Rbf.

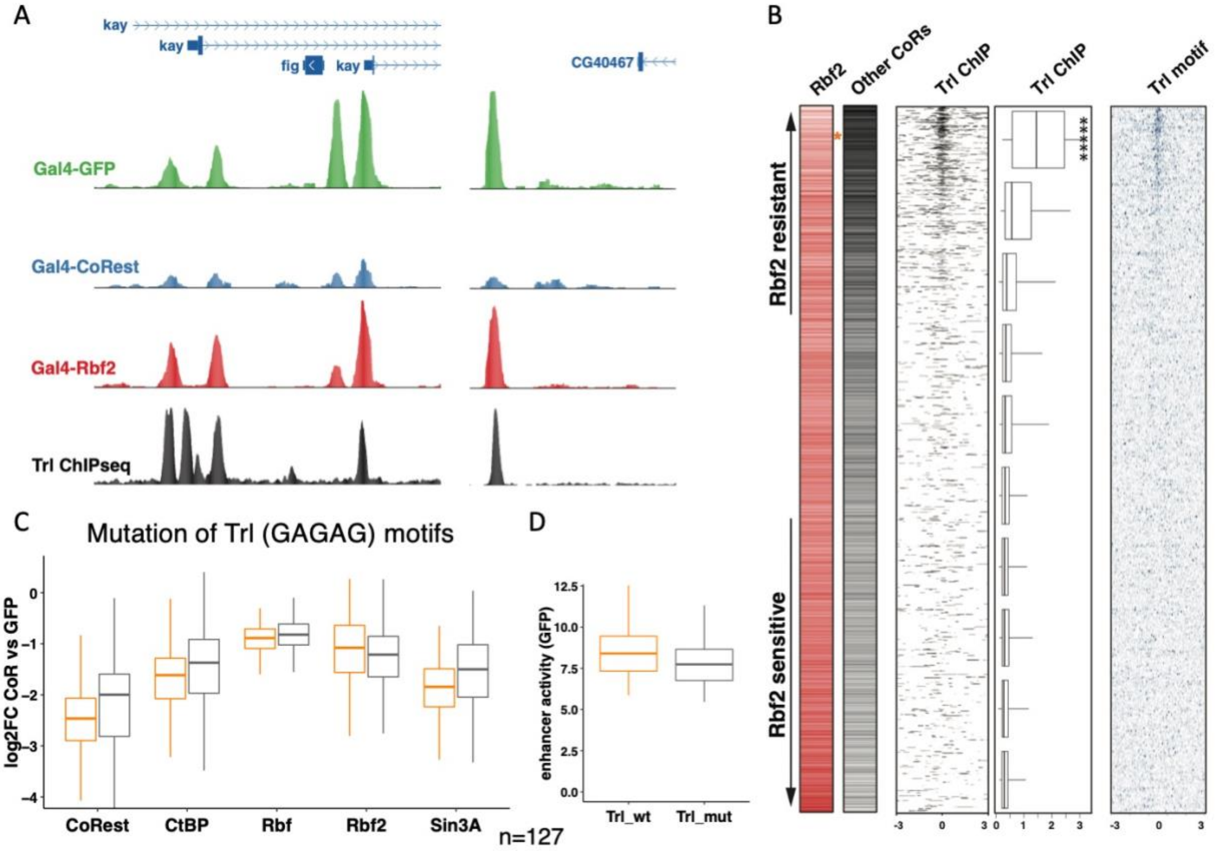


**Fig. S4. Trl confers resistance specifically towards Rbf2 repression.**
(**A**) USCS screenshots visualizing the repression by CoRest and Rbf2 of Trl bound and unbound enhancers. (B) Ranking of all enhancers from the gw-screen, based on their sensitivity towards Rbf2 repression versus towards their sensitivity towards the other 4 CoRs. Rbf2 specific resistant enhancers are on the top. Trl ChIP-seq signal is plotted in a heatmap and quantified in boxplots, Trl motifs are plotted in a heatmap. (**C**) Repression (log2FC enhancer activity CoR vs GFP) for each CoR in the screen, wt enhancers (orange) and their counterparts in which Trl motifs have been mutated (black). (**D**) Spike-in normalized enhancer activity (Gal4-GFP) in log2 scale.

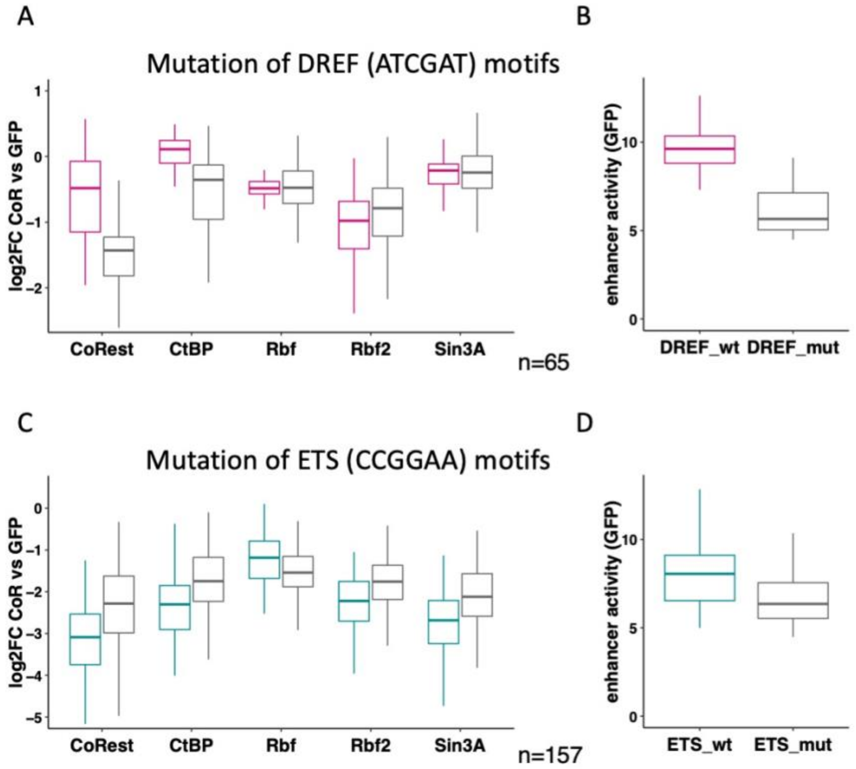


**Fig. S5. Motif mutations effect on enhancer activity and repression by all CoRs**
(**A,C**) Repression (log2FC enhancer activity CoR vs GFP) for each CoR in the screen, wt enhancers (color) and their counterparts in which the respective motifs have been mutated (black). (**B,D**) Spike-in normalized enhancer activity (Gal4-GFP) in log2 scale.

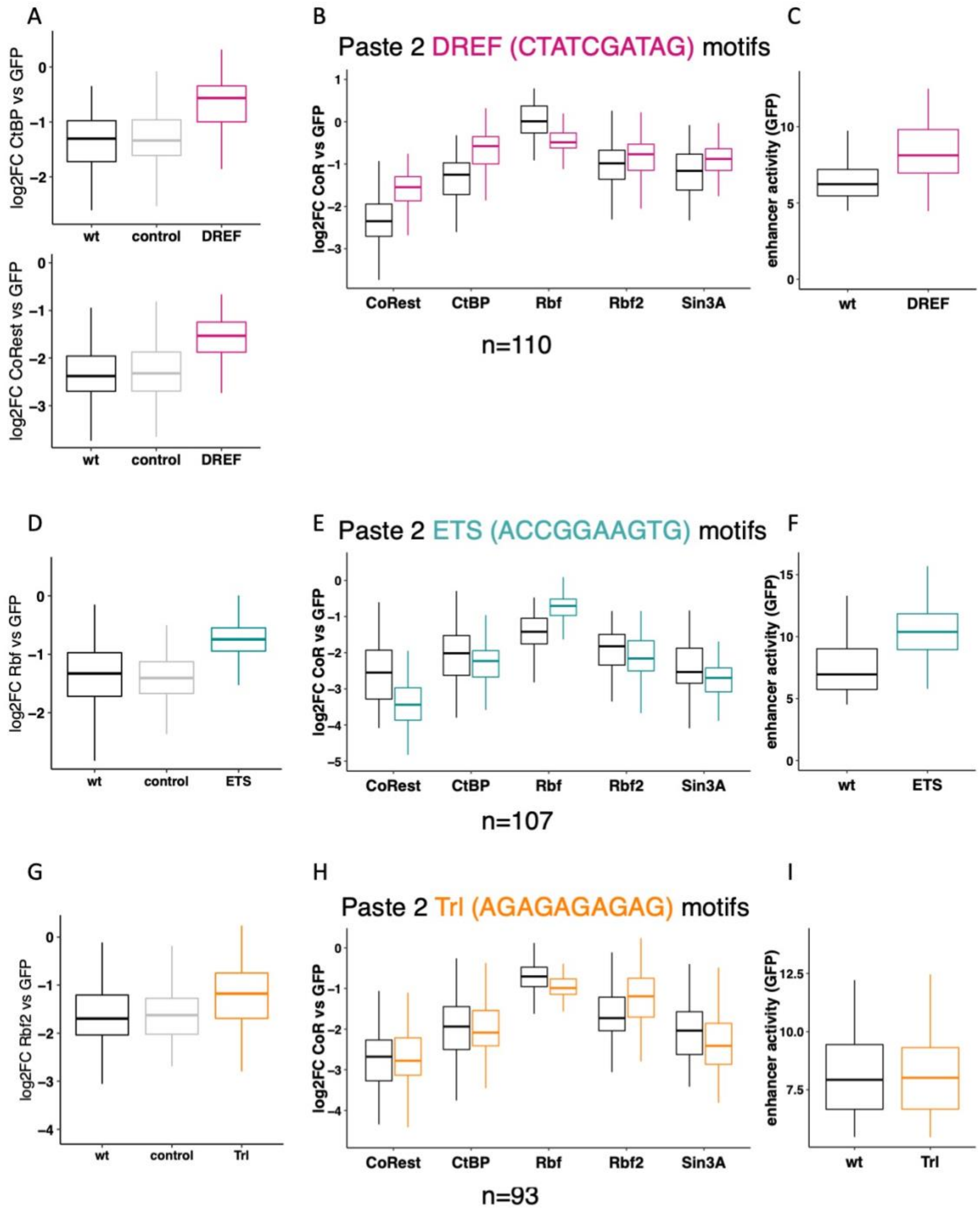


**Fig. S6. Motif addition effects on enhancer activity and repression by all CoRs**
(A,D,G) Repression (log2FC enhancer activity CoR vs GFP) of wild type enhancers (black), enhancers when the control motifs (ACE2 and FOX) are added (grey) and when the resistant motif is added (color). (B,E,H) Repression (log2FC enhancer activity CoR vs GFP) for each CoR in the screen, wt enhancers (black) and with respective motif pasted (color). (C,F,I) Spike-in normalized enhancer activity (Gal4-GFP) in log2 scale.
